# Mapping pontocerebellar connectivity with diffusion MRI

**DOI:** 10.1101/2021.09.17.460781

**Authors:** Paul-Noel Rousseau, M. Mallar Chakravarty, Christopher J. Steele

**Affiliations:** Department of Psychology, Concordia University, Montreal, QC, Canada; Cerebral Imaging Center, Douglas Mental Health University Institute, McGill University, Montreal, QC, Canada; Department of Neurology, Max Planck Institute for Human Cognitive and Brain Sciences, Leipzig, Germany

**Keywords:** Cerebellum, Pons, Pontocerebellar, Connectivity, Diffusion MRI

## Abstract

The cerebellum’s involvement in cognitive, affective and motor functions is mediated by connections to different regions of the cerebral cortex. A distinctive feature of cortico-cerebellar loops that has been demonstrated in the animal work is a topographic organization that is preserved across its different components. Here we used tractography derived from diffusion imaging data to characterize the connections between the pons and the individual lobules of the cerebellum, and generate a classification of the pons based on its pattern of connectivity. We identified a rostral to caudal gradient in the pons, similar to that observed in the animal work, such that rostral regions were preferentially connected to cerebellar lobules involved in non-motor, and caudal regions with motor regions. These findings advance our fundamental understanding of the cerebellum, and the classifications we generated provide context for future research into the pontocerebellar tract’s involvement in health and disease.

## Introduction

The cerebellum, a structure containing approximately as many neurons as the whole cerebral cortex, has historically been considered as being chiefly involved with motor processes. Recent advances have led to a paradigm shift in our conceptualization of the cerebellum, and we are just beginning to appreciate its involvement in a breadth of higher cognitive processes (Schmahmann, 2019; Strick et al., 2009). Unlike the cortex, the cellular architecture of the cerebellum is highly invariant, this implies that it is performing similar operations albeit on different inputs with different outputs (Schmahmann, 2019). An understanding of the structural connectivity of the cerebellum is therefore essential to understanding its contribution to different cognitive processes and to the effects of cerebellar syndromes. Damage to different parts of the cerebellum and associated white matter pathways will produce specific patterns of impairments in motor and non-motor functioning (Rapoport et al., 2000). While the white-matter connectivity of the cerebellum has been characterized with tract-tracing studies in non-human primates (Brodal, 1979; Schmahmann, Rosene, et al., 2004; Schmahmann & Pandya, 1997), our knowledge of the more fine-grained aspects of its connectivity in humans is still limited. Diffusion weighted imaging, particularly with recent methodological advancements provides an opportunity to investigate the white-matter connectivity of the cerebellum *in vivo* (Granziera et al., 2009; Salamon et al., 2007; Steele et al., 2017a).

The cerebellar cortex receives cortical inputs via the pons and in turn projects back to the cerebral cortex predominantly via the dentate nucleus and the thalamus. These cortico-cerbellar loops are comprised of parallel subcircuits linking spatially and functional distinct areas of the cortex and cerebellum (Schmahmann et al., 2019). The cerebellar cortex is comprised of ten lobules based on its gross anatomy (Schmahmann et al., 1999): the anterior lobe (lobules I through V and parts of VI) in addition to lobule VIII receive projections primarily from motor regions of the cortex, in addition to projections from the spinal cord. Conversely, the remainder of lobule VI Lobules VIIA and VIIB, crus I and II are connected to association areas of the cortex (Schmahmann, 2019). Previous work has established that both the pons and the dentate nucleus exhibit topographic patterns of organization based on their afferent connections (Schmahmann & Pandya, 1997; Steele et al., 2017a). In this paper, We set out to investigate the patterns of anatomical connectivity between the pons and different cerebellar regions in humans which has not been systematically investigated until now.

Studying cortico-cerebellar connectivity with diffusion tractography presents some unique challenges. First, cortico-cerebellar connections are polysynaptic and it is unclear how effective diffusion tractography is at resolving these polysynaptic connections. There are also a large number of intersecting fibers in the cerebellum and brainstem, meaning that the diffusion tensor model (in which only one fibre orientation is assumed per voxel) is inadequate for studying white matter architecture in this region (Takahashi et al., 2013). High-angular resolution diffusion imaging (HARDI) data which allows for the definition of fibre orientation distribution functions (fODFs) with constrained spherical deconvolution improves the characterization of these fiber populations and affords new opportunities for studying cerebellar connectivity (Dell’Acqua et al., 2013; Steele et al., 2017b; Tournier et al., 2007).

Our objective with the current study was to address the gap that currently exists in the literature and systematically characterize the white matter connectivity between the pons and lobules of the cerebellum. We first performed probabilistic tractography on the connections between the cerebellar lobules and the pons which travel through the middle cerebellar peduncle (MCP). Our goal was to characterize their spatial distribution, and subsequently generated parcellations of the pons and MCP based on this connectivity. The current findings further our basic understanding of normative patterns of cerebellar connectivity in humans. Moreover, they help us to predict with greater precision how damage to the cerebellum, and its associated white matter fibres may produce specific types of impairments in cognitive and motor function.

## Materials and Methods

### Participants

100 unrelated participants (50 females, average age = 29) were randomly selected from the Human Connectome Project open-access dataset (www.humanconnectome.org) (Glasser et al., 2016; Van Essen et al., 2012) and represented subset of data from a prior study (Steele & Chakravarty, 2018). Written informed consent was obtained from each participant and data was used in compliance with ethical guidelines of Concordia University and The Human Connectome Project. 100 subjects were selected to balance between the computational demands of the tractography with the requirement of a large and representative sample. Structural imaging data were acquired on a 3T Siemen Connectome Skyra scanner. T1w (0.7 mm iso, TI/TE/TR = 1000/2.14/2400 ms, FOV = 224×224 mm) and diffusion weighted imaging (1.25 mm iso, TE/TR = 89.5/5520 ms, FOV = 210×180 mm, multiband 3, b-values = 1000/2000/3000 s/mm2, 90 diffusion directions across each b-value).

### Image Processing

#### Cerebellar Lobular Segmentation – MAGeT Brain

Segmented cerebellar lobules were used as regions of interest for the tractography analysis. Segmentations were performed for a prior study by Steele & Chakravarty (Steele & Chakravarty, 2018) based on preprocessed T1 weighted images from the HCP using the Multiple Automatically Generated Templates segmentation tool (Chakravarty et al., 2013; Park et al., 2014). The method is described in detail in Steele & Chakravarty (Steele & Chakravarty, 2018), but to summarize: Non-linear registrations are performed between five manually labelled cerebellar lobular atlases and T1 weighted images from twenty-one participants from the dataset yielding a set of five cerebellar labels for each of the template participants. The resulting templates are then warped to each of the participants, resulting 105 cerebellar segmentations per subject. A majority vote procedure is then used to yield the final segmentation of the cerebellar lobules. The segmentations used for seeds ROIs in the connectivity analysis consisted of lobules III-VI, Crus I-II, lobule VIIb, lobule VIIIA, and lobule VIIIb.

#### Segmentation of Pons and Brainstem Structures - Freesurfer

Brainstem structures were used as regions of interest for the tractography analysis. Segmentation of brainstem structures was performed using a tool implemented by Freesurfer (v6.0) (http://surfer.nmr.mgh.harvard.edu). In brief, this tool uses a Bayesian segmentation algorithm which uses a probabilistic atlas of the brainstem and surrounding brain structures based on manually labeled scans to generate segmentations of the medulla oblongata, pons, midbrain, and superior cerebellar peduncle (SCP) based on T1w images (Iglesias et al., 2015). From these segmentations we used masks of the pons, medulla, and midbrain in the subsequent tractography analysis. In order to prepare ROIs for tractography, the individual subject pons masks were first dilated by three voxels. A subtraction was performed between this dilated mask and the undilated pons mask to produce a pons shell that was divided right and left pons shell masks. Tractography was performed from the left hemisphere cerebellar lobules, and the right pons shell mask served as an exclusion mask in that analysis. Streamlines passing through this area were excluded. The medulla oblongata, midbrain and SCP were also similarly used as exclusion masks for the tractography.

#### DWI Preprocessing

3T diffusion weighted images were preprocessed by the standard HCP pipeline, the steps are delineated in detail by Glasser et al. (Glasser et al., 2016). In brief, these include intensity normalization, distortion estimation and correction and a gradient nonlinearity correction. A rigid body transformation is performed to register the resultant images to the T1w structural image.

#### Tractography between cerebellar lobules and pons

MRtrix3 (Tournier et al., 2019) was used for the estimation of the fiber orientation distribution function (fODF) which was derived with constrained spherical convolution and to perform probabilistic tractography. To investigate whether the pons exhibits a topographic pattern of connectivity with the cerebellum similar that which has been observed in the animal literature(Biswas et al., 2019; Schmahmann & Pandya, 1997), we performed tractography between lobules III, IV, V, VI, Crus I and II, VI, VIIB, and VIIIA/B in the left hemisphere of the cerebellum and the pons. The segmented lobules were individually used as seed masks, and the pons used as the target mask. The right pons shell mask, medulla oblongata, midbrain, and SCP were all used as exclusion masks. This configuration of inclusion and exclusion masks ensured that only streamlines terminating pons were preserved and that streamlines transitioning through the pons and terminating in other areas were excluded. In order to prevent the final streamline counts from being biased by individuals with larger brains, the total number of streamlines from each lobule was constrained to be proportional to the volume of the largest lobule (Steele et al., 2017a). In practice, the volume of each of lobule was computed for each participant, the number of streamlines originating from the largest lobule (Crus I in most individuals) were set to 50,000, and the number of streamlines for the remaining lobules were set as a proportion of their volume to the largest lobule. For example, if a lobule had half of the volume compared to the largest lobule, streamlines would be set to 25,000. The result of this procedure was that while each lobule projects a different number of streamlines, they each project the same number of streamlines per voxel. Subsequent to tractography, individual tracts were converted to streamline count images which were used in the group template formation and the final segmentations.

#### Group Co-registration

In order to put the individual subject data into the same space, a group template based on the streamline count data was generated using the ANTs registration software (Avants et al., 2011). Total streamline count images across individuals (the sum of the individual streamline count images for each of the lobules) were non-linearly co-registered to one another and averaged to form a group template (antsMultivariateTemplateConstruction2, demons similarity metric). Individual native-space streamline count data for the individual lobules were then warped to the final group template.

#### Lobule-specific Visitation Maps

In order to visualize the distribution of streamlines within the pons and MCP, individual streamline counts for each lobule were scaled by the number of streamlines from the lobule to generate comparable proportional visitation maps for each lobule. This scaling step allowed us to better account for the distribution of connectivity from the smaller lobules, and these maps served as the input for the streamline-based segmentation described in the next section. The scaled maps were then summed across individuals and normed between 0 and 1 to generate group probability maps of streamline distributions for each lobule. Intensities were then normalized according to the maximum in a slice of interest at the midline for ease of visualization.

#### Streamline Based Parcellation of the Pons and MCP

Based on the lobule specific streamline count maps derived from the tractography analysis, subject-level and group-level classifications of the pons and MCP were performed using a modified majority-vote procedure. A standard majority vote procedure, where a voxel is labeled according to the lobule which contributes the most streamlines, has the potential to under-represent the contributions smaller lobules to the final parcellation. In order to better characterize the spatial distribution of connectivity from smaller lobules of the cerebellum, we used a modified procedure where the individual lobule specific streamline count images for each subject were divided by the number of streamlines in that lobule. (relative strength of connectivity from each lobule). For purposes of comparison, we present the results parcellation results from both the scaled and non-scaled streamline data.

A parallel set of analyses were conducted on the group and subject level data. Using the lobule specific streamline count maps as inputs, voxels were labeled according to lobule map with the highest intensity in that position (in the lobule scaled approach), and according to the lobule contributing the most streamlines (in the simple majority vote approach). In order to estimate the consistency of the group labels across different participants we computed a cross-subject consistency metric. At each voxel in the final group segmentations, we computed the proportion of individual subject segmentations with the same label at that position. All summary maps and parcellations will be made available on NeuroVault (https://neurovault.org/) at the time of publication.

## Results

### Lobule Specific Tractography between the Cerebellar Cortex and Pons

Tracking was performed with the lobules as seeds and the pons as the target, and though axonal projections originate in the pons and terminate in the lobules, for simplicity of description we describe connectivity from the point of view of the lobules. Streamlines originating in the cerebellar lobules change direction along the superior-inferior and lateral-medial axes in their course towards the pons. Lobule specific tractography from each of the lobules in a representative participant is depicted in Figure 1. Streamlines from within lobules III, IV, V travel inferiorly, turn and travel anteriorly (with lateral extents matching their exit from the lobules) as they enter the region of white matter surrounding the dentate nucleus and eventually form the middle cerebellar peduncle. In contrast to the more anterior lobules, streamlines from lobules VI, Crus-I, and Crus-II follow a more direct and linear track as they join the white matter of the cerebellum. The majority of streamlines from VI and Crus I (and all lobules anterior to them) pass over the superior surface of the dentate nucleus, while streamlines from Crus-II bifurcate to travel above and below the dentate nucleus. Streamlines from lobules VIIB, VIIIA, and VIIIB travel in the superior direction, converge, and turn to continue anteriorly. Streamlines from lobule VIIB bifurcate around the top and bottom of the dentate nucleus, and streamlines from VIIIA and VIIIB largely travel inferior to the dentate nucleus. Streamlines from each of the lobules make up the single bundle of the MCP immediately posterior to the pons that turns to travel medially as it enters the pons. The streamlines then bifurcate along the anterior-posterior axis to form two bundles anterior and posterior to the corticospinal tract. As streamlines enter the pons they also fan out in the rostral-caudal axis, with each lobule showing different patterns of distribution along these axes. In general, streamlines from lobules III, IV, and V had the most prominent bifurcations that form distinct rostral and caudal bundles.

**Figure 1.**
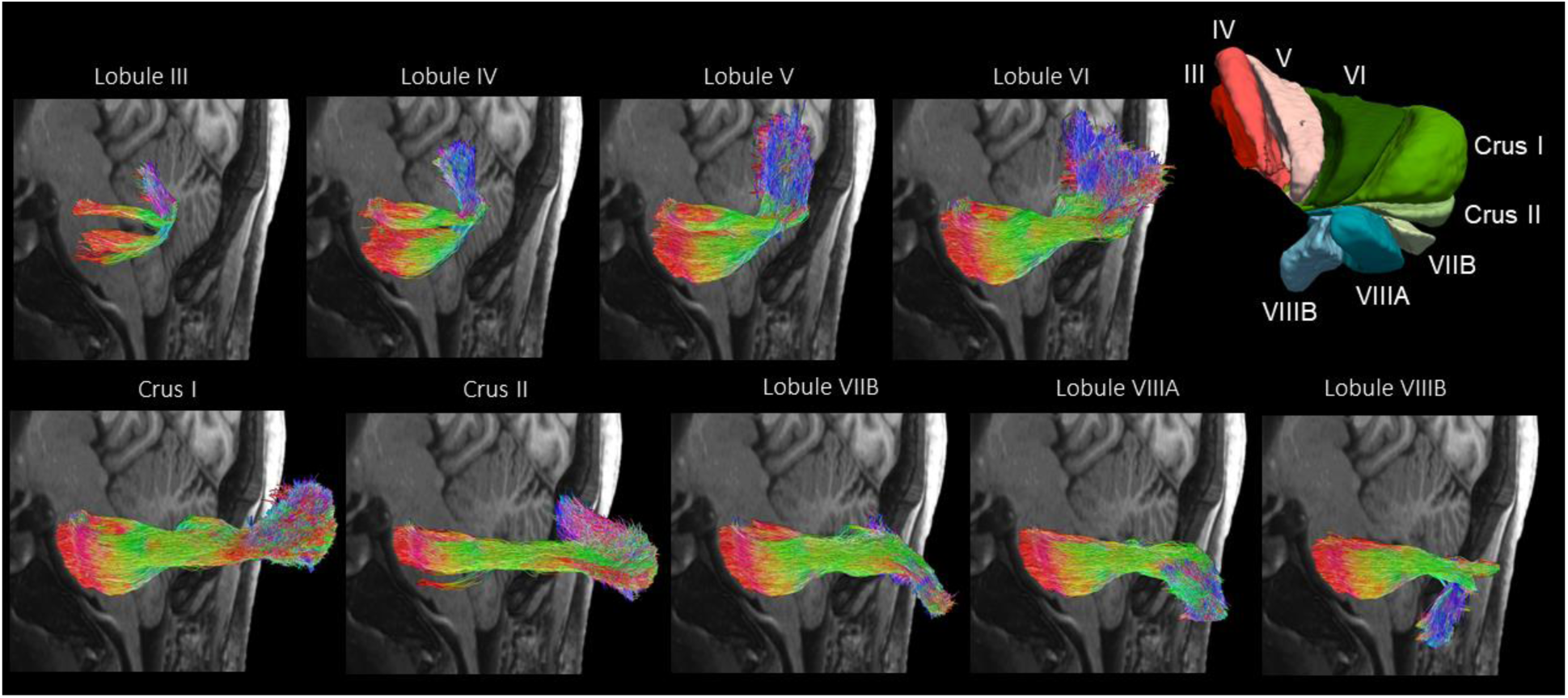
Tractography results in single subject for the individual lobules showing their trajectory displayed at the midline of the brain at an oblique angle. The color of the streamlines indicates their directionality: blue is superior/inferior, red is medial/lateral, and green is anterior/posterior. The graphic in the top right is a 3D view of the cerebellar lobule segmentation shown at an oblique angle. Outlier streamlines were filtered for clarity of display using scil_outlier_rejection.py from the Sherbrooke Connectivity Imaging Lab toolbox in Python (Scilpy).

### Group Level Visitation Maps

#### Pons

Figure 2 depicts group average maps of normalized streamline distributions from each of the cerebellar lobules in a sagittal cross section at the midline of the pons. The dark area in the center of each cross section represents the location of the corticospinal tract. Streamlines from lobule III were distributed in two distinct areas in the rostral and caudal aspects of the pons. Lobule IV streamlines were primarily concentrated in the inferior half, whereas lobule V streamlines were more distributed in both rostral and caudal parts of the pons. Streamlines from lobules VI, Crus and Crus II were mostly distributed in the rostral half of the pons. Lobule VIIB and VIIIA streamlines projected predominantly to more central parts of the pons and VIIIB streamlines were distributed more caudally. To get an idea whether motor and non-motor cerebellar lobules showed differential patterns of connectivity with the pons, streamlines from the putative motor (III, IV, V, VIIIA, VIIIB) and non-motor (VI, Crus I, Crus II, VIIB) lobules were summed. Streamlines from motor lobules were distributed more in the caudal half of the pons whereas streamlines from non-motor lobules were concentrated in the more rostral aspect.

**Figure 2.**
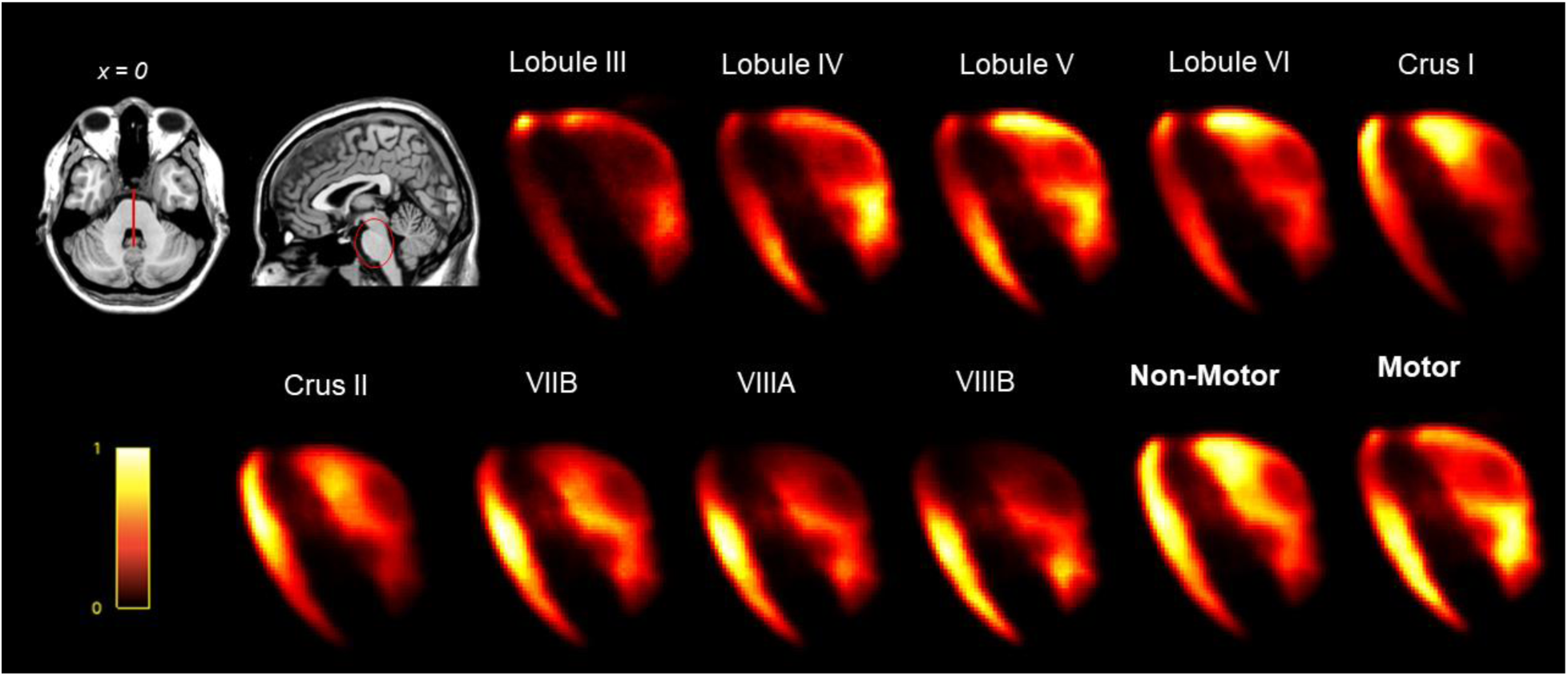
Normalized streamline distribution maps at the midline of the pons for each of the cerebellar lobules, and the sum streamlines of non-motor (VI, Crus I, Crus II, VIIB) and motor (III, IV, V, VIIIA, VIIIB) lobules. Brighter colors indicate higher density of streamlines. The dark region in the middle of each pons cross-section corresponds to the cortico-spinal tract.

#### MCP

Figure 3 depicts group average streamline distribution maps in a cross section of the MCP as it exits the cerebellum. Streamlines from lobules III, IV and V and VI are concentrated in two distinct clusters in superior and inferior portions of the medial MCP. This reflected the bifurcations for the streamlines from these lobules described above. For Crus I, Crus II, lobules VIIB, VIIIA, and VIIIB the predominant location of streamlines are organized in a clockwise fashion around the cross-sectional center of the MCP: ranging from Crus I’s concentration in the superior and lateral portion of the MCP to lobule VIIIB in the inferior and medial portion of the MCP. When streamlines from non-motor lobules (VI, Crus I, Crus II, and VIIB) were summed, they were found to concentrate in the lateral and superior quadrant of the MCP. In contrast, the summed motor lobules (III, IV, V, VIIIA, VIIIB) were clustered primarily in the inferior and medial quadrant of the MCP and to a lesser extend in the superior and medial quadrant.

**Figure 3.**
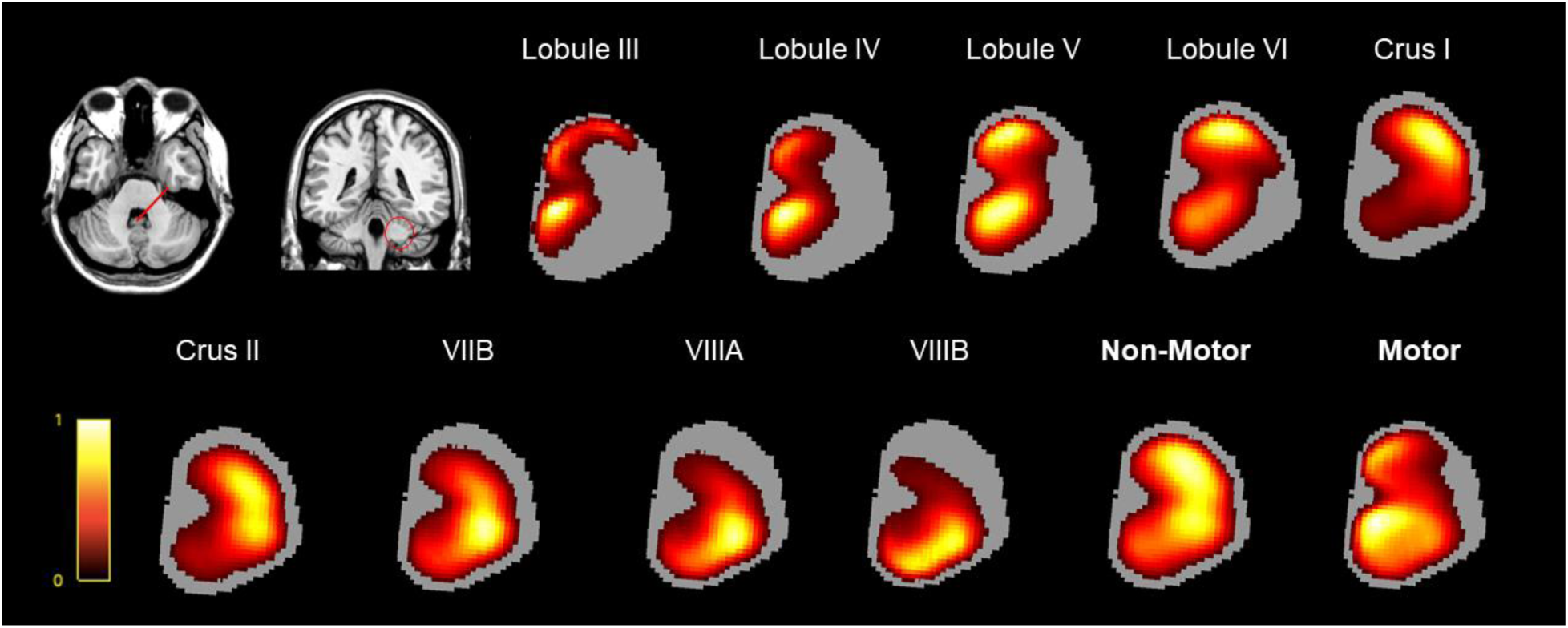
Normalized streamline distribution maps in a cross section of the MCP for each of the cerebellar lobules, and the sum streamlines of non-motor (VI, Crus I, Crus II, VIIB) and motor (III, IV, V, VIIIA, VIIIB) lobules. Brighter colors indicate higher density of streamlines.

### Lobule Scaled Majority Vote Classification

#### Group Level

##### Pons

Figure 4 depicts the results of the group-level majority vote segmentation in the pons and MCP as well as the inter-subject agreement metric using the scaled streamline count data. In the ventral half of the pons, going from rostral to caudal, we note layers corresponding to lobule III, followed by Crus I, Crus II, lobule VIIIA. Lobule VIIIB is represented in much of the caudal half of the pons. In the dorsal portion of the pons, we note small lobule V and VI areas, and more inferiorly large lobule III and IV areas. The overall spatial organization of the segmentation is similar to the organization of the cerebellar lobules in the cortex, with neighbors in the cortex generally found adjacent to each other in the pons. Lobules III and IV are notable exceptions, appearing at multiple different locations in the pons. Inter-subject agreement is highest in the periphery of the pons cross-section, in particular in the area that corresponds to Crus I. Consistency of the labels is lowest in the center of the pons where there is a greater overlap of streamlines belonging to different lobules.

**Figure 4.**
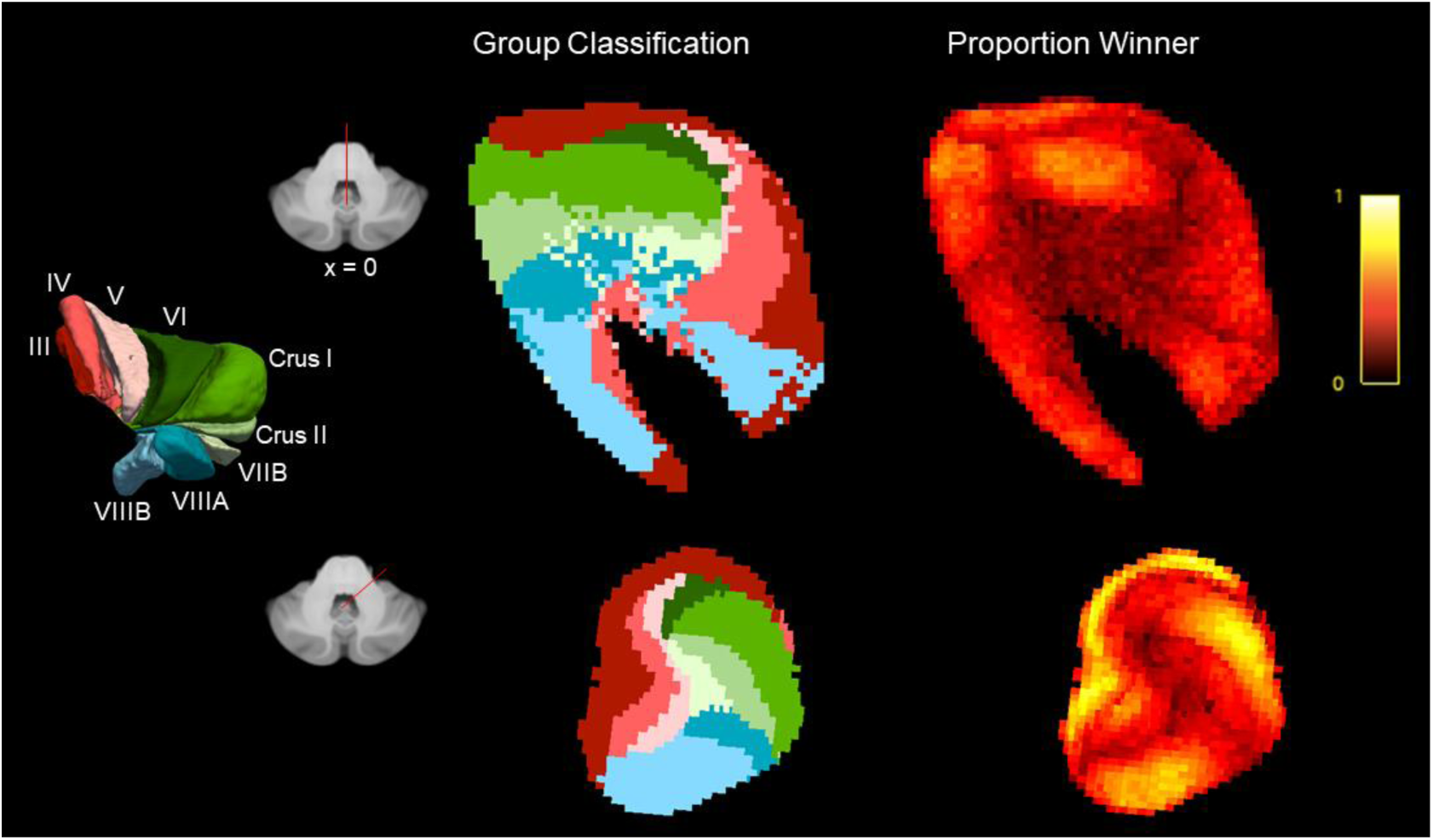
On the left are results of the majority vote classification using the scaled streamline count data for the pons (top) and MCP (bottom). On the right are the intersubject agreement maps for the pons and MCP, here brighter colours indicate higher consistency of the label across subjects.

##### MCP

In the MCP, lobule III is represented in the most medial periphery of the cross section, followed by layered medial to lateral stacking of lobules IV, V and VI towards the center of the MCP. In contrast, Crus I, Crus II, Lobules VIIA and VIIB are layered in more of a superior to interior fashion in the lateral half of the cross section. There is a small area corresponding to lobule VIIB at the very center of the cross section. Inter-subject agreement is highest in the periphery, especially in the regions of lobules III, Crus I and VIIB. As with the pons, consistency is lowest in the center where there is more overlapping of streamlines.

#### Individual Participant Level

##### Pons

The results of the subject level parcellation in the pons using the scaled streamline data is presented for 8 representative participants (4 females) in figure 5. Although there is heterogeneity in the exact position and size of specific labels, there are a number of consistent observations between individuals and with the group segmentation. Labels show a similar layering pattern which, for the most part, recapitulates the spatial configuration of the cerebellar lobules.

**Figure 5.**
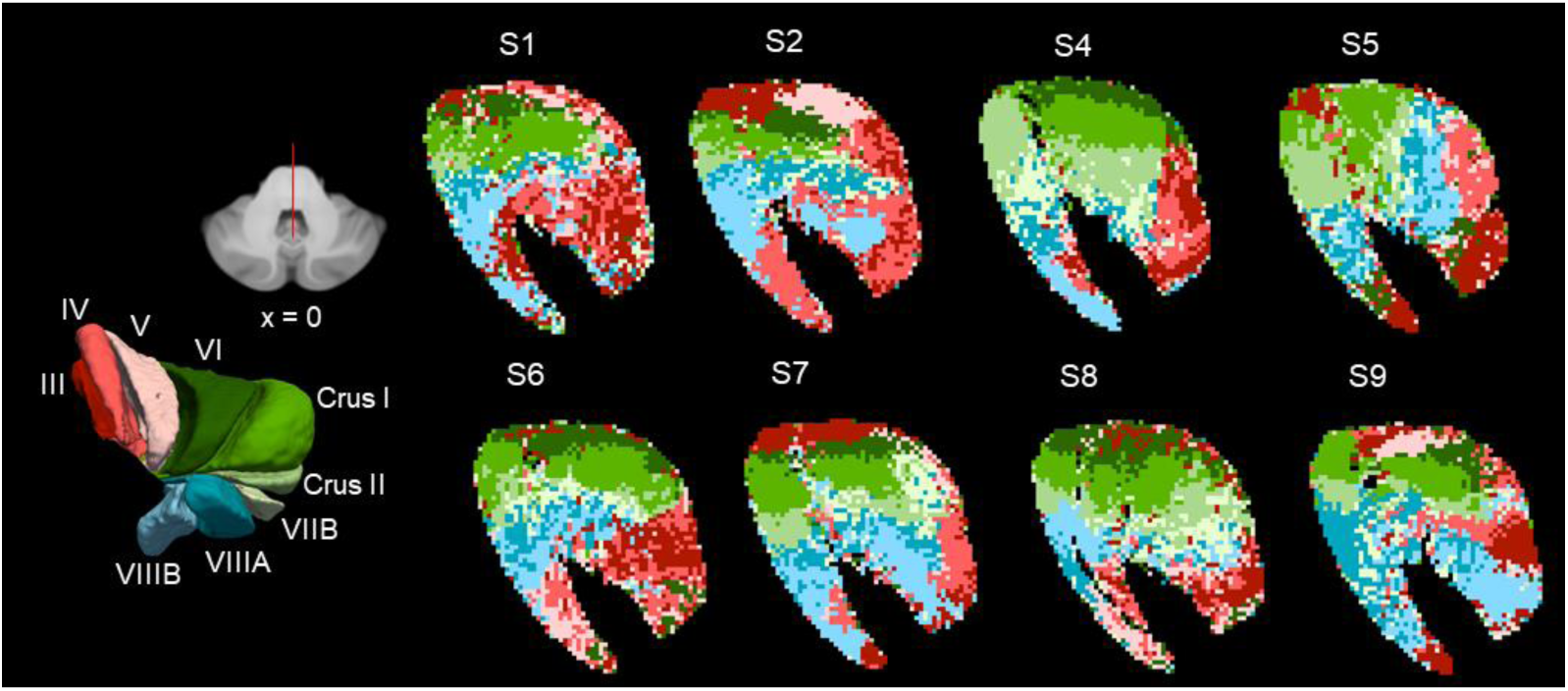
Results of the majority vote segmentation of the pons using the scaled streamline count data in 8 representative participants.

##### MCP

In the MCP, depicted in figure 6, we observe consistency in the general spatial configuration of the individual parcels, with some variability in their exact position and size.

**Figure 6.**
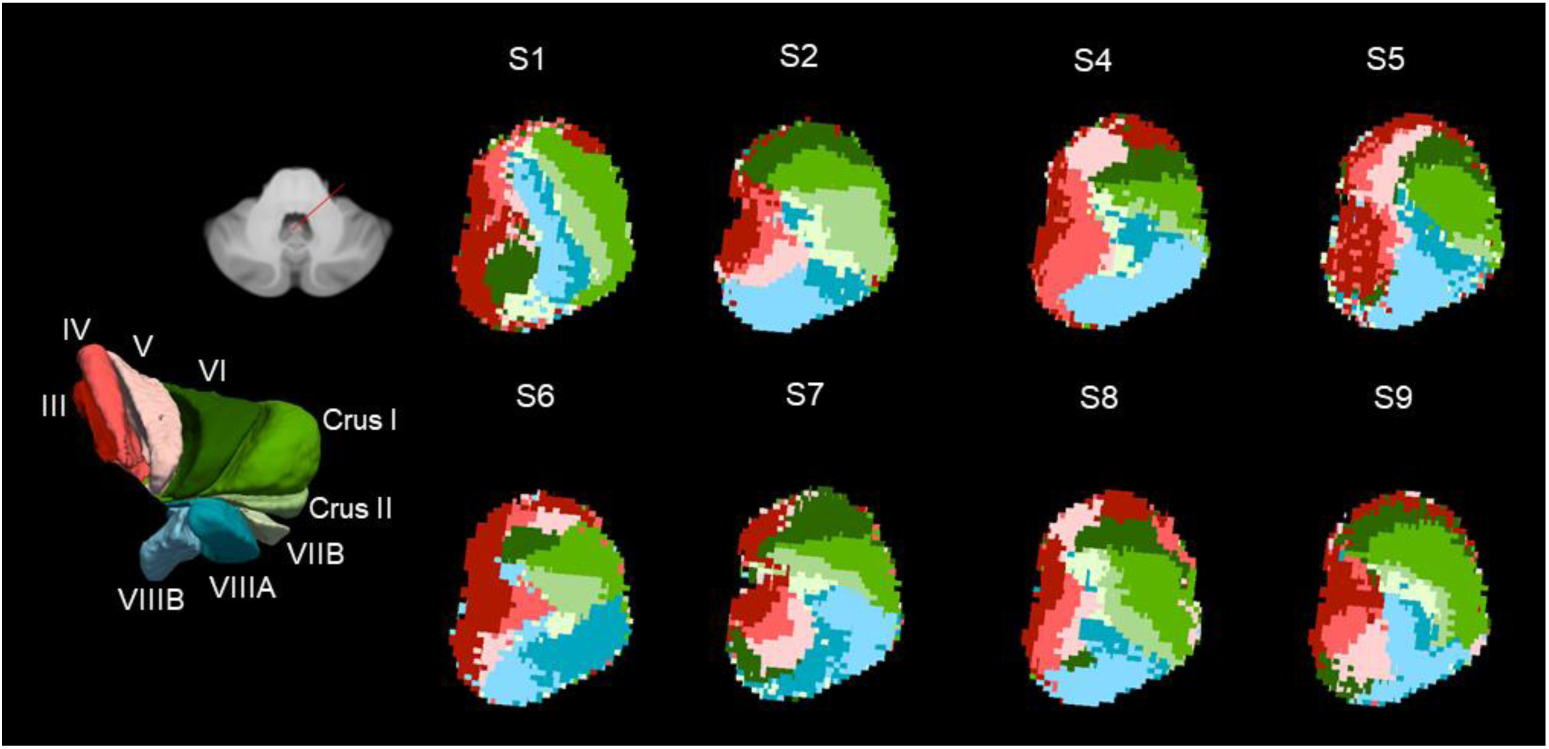
Results of the majority vote segmentation of the MCP using the scaled streamline count data in 8 representative participants. Shown in a cross-section of the MCP located proximal to the pons.

Similar to the group segmentation: medially, we observed areas corresponding to lobule III, IV and V. In the lateral half of the MCP cross section, going from superior to inferior, we observed layers corresponding to Crus I, Crus II, lobules VIIIA and VIIIB as identified in the group map. However, we also noted that there was variability in the orientation of the layers, with some participants exhibiting a more medial to lateral organization.

### Simple Majority Vote

#### Group Level

##### Pons

Presented in figure 7 are the results for the group-level majority vote in a cross-section of the pons and MCP, as well as the inter-subject agreement metric using the non-scaled data. The segmentation is generally consistent with that performed with the scaled data, but as expected it only includes the larger lobules of the cerebellum. In order of size, there are areas corresponding to lobules VI, Crus I, Crus II and VIIB. The lobule VI area starts in the dorsal part of the rostral pons, and then wraps around to cover much of the caudal half the pons. The Crus I area is in the rostral half of the pons, and the Crus II and VIIB areas are immediately caudal to this and are focused in the ventral portion. The agreement metric shows that the Crus I label and Lobule VI labels were broadly consistent across participants, in the rostral half of the pons. The Crus II and lobule VIIB labels were less consistent across participants.

**Figure 7.**
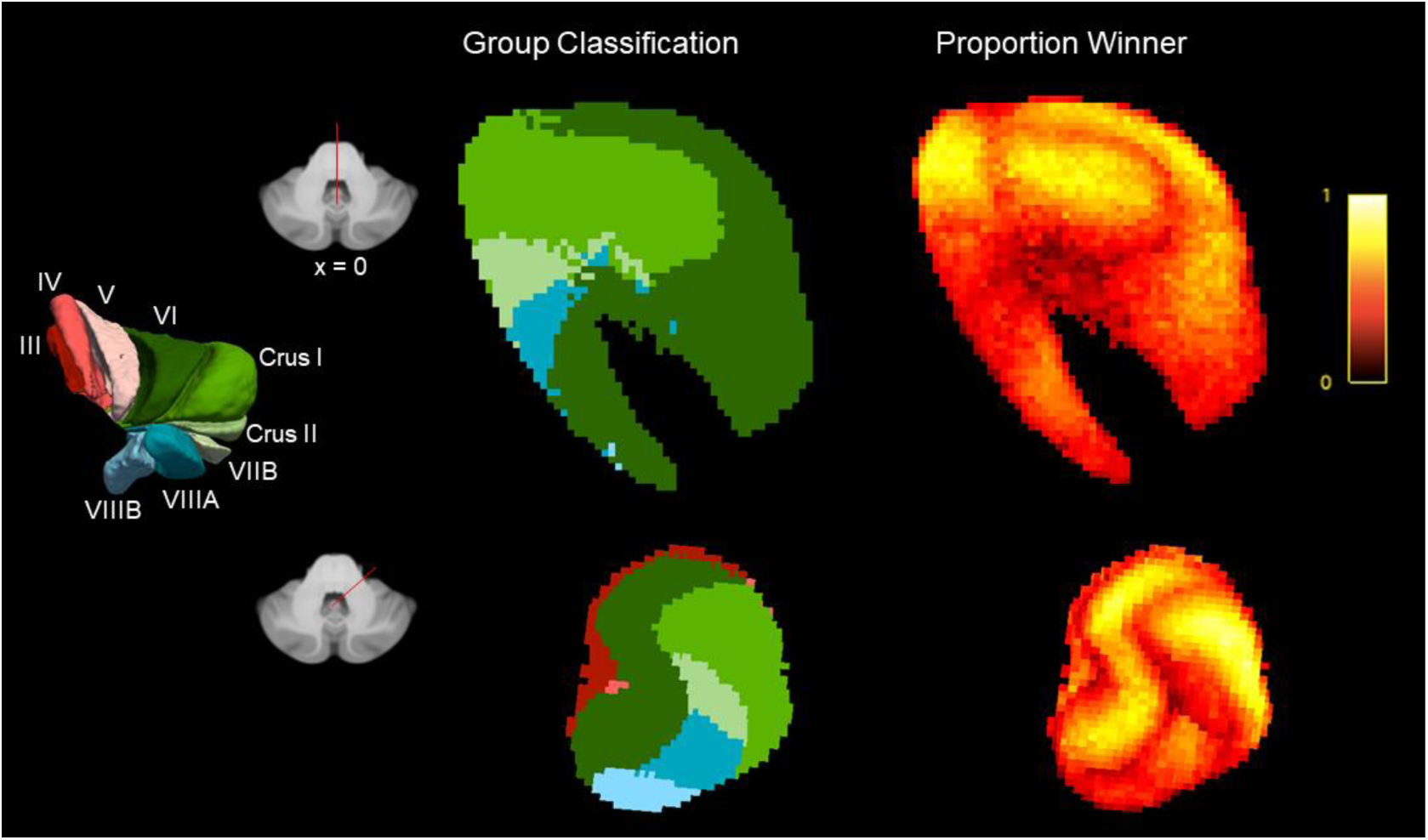
On the left are results of the majority vote classification using the raw streamline count data for the pons (top) and MCP (bottom). On the right are the intersubject agreement maps for the pons and MCP, here brighter colours indicate higher consistency of the label across subjects.

##### MCP

At the level of the MCP, the majority vote results again parallel those from the scaled analysis. Again, however, we note that the larger lobules are overrepresented in the segmentation. In a cross-section most proximal to the body of the pons, we note a small area in the most lateral portion corresponding to lobule III and a large area in the lateral half corresponding to Lobule VI. Going clockwise we find smaller areas corresponding to Crus I, Crus II, VIIIA, and VIIIB. With the agreement metric, similar to with the pons, we note high consistency of the Lobule VI and Crus I labels, but considerably more heterogeneity for the other labels.

#### Individual Participant Level

##### Pons

The results of the subject-level pons segmentations for the same 8 participants using the non-scaled majority vote data are depicted in Figure 8. Similar to the group analyses, the segmentation is dominated by a small number of lobules. As with the scaled analysis, there is considerable heterogeneity across individuals, but some consistent patterns - the areas corresponding to Crus I and Crus II streamlines occur in the rostral half of the pons, whereas the area corresponding lobule VI streamlines occur in most rostral and caudal parts of the pons. This likely reflects the bifurcation noted in the tractography and two distinct clusters of streamlines noted in the average streamline distributions in the MCP cross-section.

**Figure 8.**
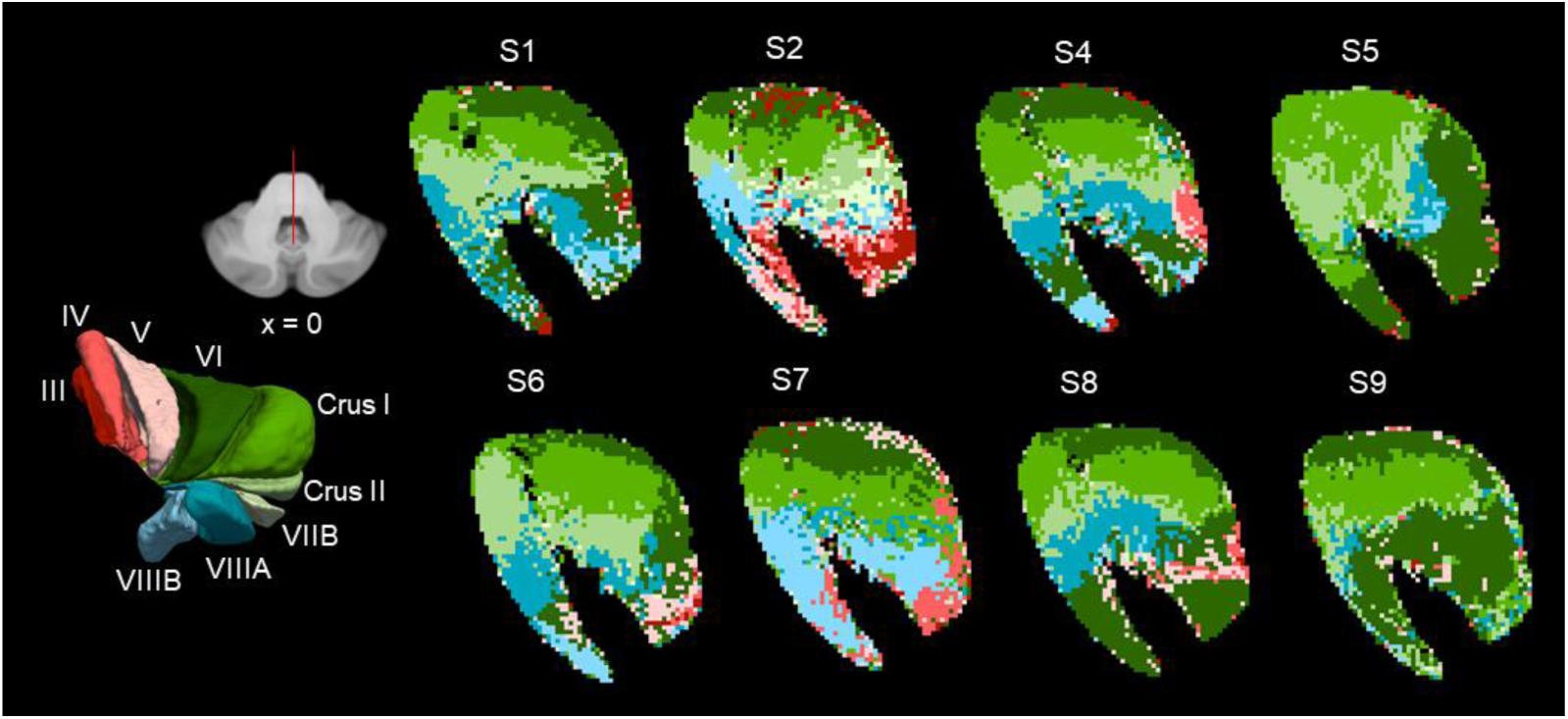
Results of the majority vote segmentation of the pons using the raw streamline count data in 8 representative participants. Shown the midline of the brain (x = 0).

##### MCP

Figure 9 depicts the results for the majority voting procedure in the same 8 participants in a cross section of the MCP. In most participants, we observe a small area corresponding to lobule III in the most medial portion of the cross-section. We then note a large area in the medial half corresponding to lobule VI. Crus I, Crus II, VIIIA, VIIIB are organized in a layered fashion in the lateral half of the pons. As with the scaled data, these can be layered on more of a medial to lateral axis in some participants.

**Figure 9.**
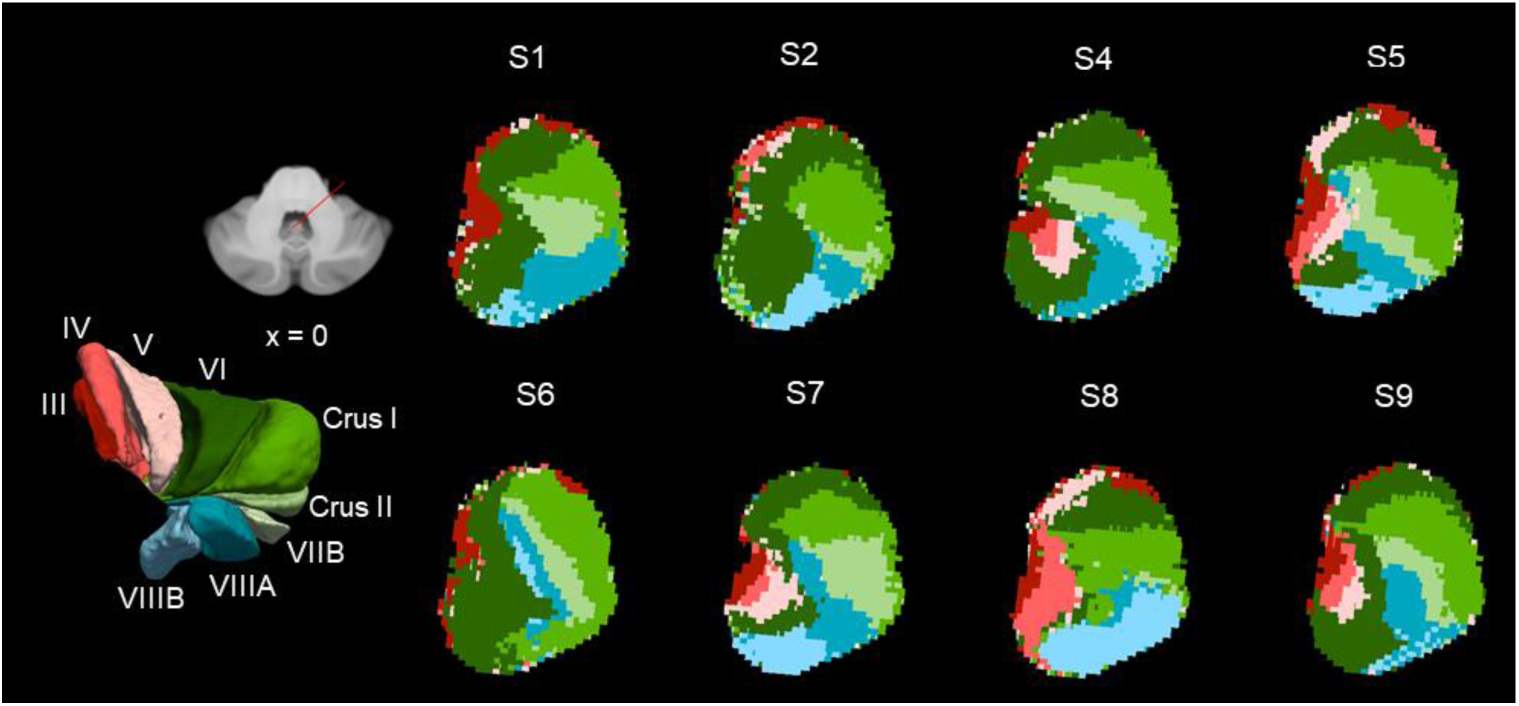
Results of the majority vote segmentation of the MCP using the raw streamline count data in 8 representative participants. Shown in a cross-section of the MCP located proximal to the pons.

## Discussion

We used *in vivo* diffusion imaging to reconstruct pontocerebellar projections and demonstrated segregated patterns of connectivity in the MCP and between the pons and cerebellar cortex. Consistent with animal anatomical work, we find a distinct pattern of connectivity such that the motor lobules of the cerebellum are more highly connected to the rostral pons and non-motor lobules are more connected to caudal pons. Our observations parallels observations from the stroke data in humans which shows different patterns of behavioural disturbances depending on lesion location in the pons. Here it is important to note that cerebellar lobules cannot be summarily classified as being motor or non-motor, this somewhat artificial distinction does allow for us to situate our findings in regards to previous literature. To our knowledge, our study is also the first to classify the pons and MCP based on their connectivity to the cerebellar lobules. Our novel approach yielded a classification of the pons and MCP which provides further evidence for the topographic organization in these structures which are likely to mediate the cerebellum’s involvement in behaviour, cognition, and emotion. The classification also provides future researchers with a framework for disentangling the contributions of different regions of the MCP and pons and MCP to these different functions.

### Spatial Configuration of the MCP

The spatial configuration and trajectory of the MCP has previously been described using dissections in a small sample of human brains (Akakin et al., 2014; Jamieson, 1910; Perrini et al., 2013). Our lobule specific tractography approach recapitulated the prominent organizational principles noted in these studies. In one of the earliest published studies to describe the trajectory fibres in the MCP, Jamieson (Jamieson, 1910) noted the existence of two distinct white matter bundles in the pons: a more superficial bundle in the rostral pons traveling to the superior posterior lobe, and a deeper bundle in the caudal pons traveling to the superior lobe. We noted a similar overall pattern in our results: the rostral pons was more highly connected to the posterior lobe (primarily Crus I and Crus II), whereas the caudal pons was connected to the superior lobule. Furthermore, streamlines connecting the pons to the posterior lobe tended to be concentrated in the lateral portion of the MCP, while those connecting to the superior lobe were concentrated in the medial MCP. A similar distinction between deep and superficial bundles of the MCP was also made by Akakin et al. (Akakin et al., 2014) who studied its anatomy in relation to the dentate nucleus. They observed one bundle with a more lateral course (projecting to the posterior lobe of the cerebellum) and another travelling along a more medial course parallel to the midline. For the later bundle, the authors did not specify where its fibres terminated in the cerebellar cortex. The first observation is in line with our findings concerning the distribution of streamlines in the MCP, where streamlines connecting the pons to lobules in the posterior lobe were more concentrated in its lateral aspect. Taken together, these dissection studies lend validity to our tractography analysis in that the larger scale aspects of connectional anatomy of this brain region are reflected in our results. We did note a bifurcation in the bundles corresponding to lobules III, IV, and I this is likely due to the intersection between these and other bundles such as those originating in Crus I and II and reflects a limitation of tractography at this resolution.

### Topographic Organization of the Pons

We demonstrated that connections between the pons and the cerebellum followed a rostral-caudal gradient, with the rostral pons being more highly connected to lobules more involved with higher order and the caudal pons with areas more involved with motor control. This is likely to reflect segregation that occurs within the cerebral cortical inputs to the pons. Based on anterograde tracing work done in animals, corticopontine projections appear to be characterized by patterns of divergence and segregation. Small cortical areas tend to project throughout the rostro-caudal extend of the pons, and to distinct regions which form patchy clusters with minimal overlap (Brodal & Bjaalie, 1997). Keeping in mind the fractured topology that appears in these projections and terminations in the pons, we note some larger organizational principles and their relevance to the present study. Anterograde tracing studies in macaques found that corticopontine projections from primary motor cortex terminate primarily in the caudal half of the pons, whereas projections from associative cortical areas are distributed throughout the pons but primarily in the rostral half (Schmahmann, Rosene, et al., 2004; Schmahmann & Pandya, 1997). Earlier lesions studies, also in macaques, found a similar overall pattern of a superior to inferior gradient in non-motor and motor cortical inputs to the pons (Brodal, 1978; Wiesendanger et al., 1979). While our findings are different in the respect that they demonstrated a rostro-caudal gradient in terms of pontocerebellar connections, based on tracing work done in rodents and macaques, we expect these to show spatial correspondence to the corticopontine projections at the gross level (Biswas et al., 2019; Brodal, 1982).

Pontocerebellar projections have been less extensively researched, but the existing literature is consistent with the notion of distinct regions of the pons projecting to motor and non-motor areas of the cerebellar cortex. Recent task-based and resting state functional imaging studies have demonstrated functionally distinct areas in the cerebellum (Buckner et al., 2011; King et al., 2019), it follows that white-matter input to the cerebellum from the pons should follow a similar level of segregation. Studies using horseradish peroxidase retrograde tracing found that the anterior lobe of the cerebellum receives fibres primarily from the caudal pons, whereas the posterior lobule received fibres mostly from the rostral part of the pons (Brodal, 1979, 1982). More recently, Biswas and colleagues (2019) reconstructed the trajectory of individual pontocerebellar axons in mice. Notably, they observed a core-shell organization wherein the central part of the pons and the surrounding areas projected to specific zones within the cerebellum. While they found that single axons diverged into multiple collaterals these tended to innervate particular combinations of lobules. These patterns of divergence may account for how a single motor representation in the caudal pons gives rise to a multiple motor representations in the anterior lobe and inferior portion of the posterior lobe of the cerebellum. In fact, similar to our observations, they found that the caudal pons projected both to lobules II-V as well as lobule VIII. This provides further context for our observation that inputs from to motor and non-motor areas of the cerebellum originated in spatially distinct areas of the rostral and caudal pons, and that adjacent lobules tended to connect to similar areas in the pons.

In humans, the rostral to caudal functional gradient in terms of non-motor and motor function within the pons has been demonstrated in lesion studies which have found cognitive and affective disturbances resulting from damage to the rostral part of the pons (Kim Jong S. et al., 1995; Schmahmann, Ko, et al., 2004). The impairments noted here may result from damage to cerebrocerebellar circuitry originating in higher order cortical areas. Our study offers new context to these findings in demonstrating the preponderance of connections to higher order regions of the cerebellum originating in the superior pons.

To our knowledge, structure function relationships within the MCP itself have not been explicitly studied in the previous literature. Studies have consistently found degradations in cerebellar white matter in multiple sclerosis (Wilkins, 2017), and Tobyne et al. (Tobyne et al., 2018) found white matter changes in the MCP to related to cognitive impairment in this population. Our approach to characterizing the distribution of connections from different parts of the cerebellum offers additional perspective to understanding the nature of these impairments. It generates a series of testable hypotheses relating to the type of impairments we can expect based on the location of a lesion or a specific area which shows a loss of white matter integrity.

### Lobule Specific Classification of the Pons and MCP

Diffusion tractography has been previously used to segment the thalamus and the basal ganglia based on their connectivity to the cortex (Draganski et al., 2008; Johansen-Berg et al., 2005; Traynor et al., 2010). Our study is the first to apply this technique to the segmentation of the pons based on its cerebellar connectivity. This presented some unique challenges: compared with the thalamus and the basal ganglia, where different areas of the cortex project to more spatially distinct regions of these nuclei, in the pons there is more complex pattern of organization with more convergence and divergence (Brodal & Bjaalie, 1992). In this case, a standard-majority voting approach is less valuable in that it results in a segmentation which underrepresents the smaller cerebellar lobules that project fewer streamlines. By scaling the streamline counts according to the size of the lobules in each individual, we were able to produce segmentations of the pons which better reflect the spatial configuration of the of the connections to individual lobules within the pons.

### Limitations and Future Directions

We based our tractography analysis and subsequent segmentation on cerebellar lobular subdivisions. While the functional territories of the cerebellum do not map perfectly on to the lobules, as is evidenced by a growing body of resting-state and task based functional imaging research (Buckner et al., 2011; King et al., 2019), these do provide a proven means of subdividing the cerebellum across individuals. Future research could consider more data driven methods of parcellating the cerebellum based on its white matter connectivity to the pons, such as independent component analysis (Hale et al., 2015). Other approach would involve clustering streamlines into distinct bundles (Garyfallidis et al., 2018), or using clustering algorithms based on the local orientation distribution function data (Najdenovska et al., 2018).

Further, we noted some discrepancies between our observations and our expectations based on known anatomy. In the tractography analysis, we noted that streamlines from lobules III, IV, V (motor lobules) showed a prominent bifurcation as they entered the pons (Figure 1). This was reflected in the final group classification where we found rostral and caudal parcels corresponding to these regions (Figure 4). It is likely that this is due to the close proximity and intersection between different white matter bundles originating in the posterior cerebellum. In light of this, the finding of rostral parcels corresponding to lobules III, IV, and V should be interpreted with caution.

While our study shows that fine grained anatomical details of the white matter architecture of the cerebellum can be resolved with high quality 3T data in-vivo, it our results could be further supported with higher resolution diffusion data and ex-vivo techniques. We initially explored using the 7T diffusion data available from the HCP, but the considerable signal drop-off in the inferior regions of the cerebellum made it unusable for tractography. Even using a cerebellum specific acquisition Steele et al. (Steele et al., 2017b), encountered similar issues of poor signal in the most inferior lobules of the cerebellum. While higher in-vivo MRI does offer interesting possibilities for studying the white matter of the cerebellum, future research will have to overcome this constraint. Ex-vivo diffusion and novel techniques like tractography based polarized light imaging microscopy, which offers spatial resolutions of up to 100μm, can offer new perspectives into meso-scale connectional architecture (Axer et al., 2011).

### Conclusion

The cerebellum’s involvement in a wide range of cognitive, affective and motor processes is mediated by its connectivity to the brain. Our study contributed to our understanding of its connectivity by demonstrating that the connections between the cerebellar cortex and the pons follow a topographic pattern of organization that is similar to observations from non-human primates. Specifically, we provided evidence that areas of the cerebellum involved with motor and higher order process are connected to spatially distinct areas in the pons, via different parts of the MCP. As well as contributing our fundamental understanding of the cerebellum, our description of its pattern of connectivity as well as the parcellations of the pons and MCP provide new context for studying the relationship between damage to these areas and functional impairments.

## Acknowledgements

This work was supported by scholarships from the Quebec Bioimaging Network and Fonds de Recherche du Québec – Nature et technologies to Paul-Noel Rousseau, grants from the Natural Sciences and Engineering Research Council and the Stroke Foundation of Canada to Christopher Steele. M. Mallar Chakravarty received salary support from Fonds de Recherche du Québec – Santé and research support from Canadian Institutes of Health Research, the Natural Sciences and Engineering Research Council, and McGill University’s Healthy Brains, Healthy Lives program.

